# ARTEM: a method for RNA and DNA tertiary motif identification with backbone permutations, and its example application to kink-turn-like motifs

**DOI:** 10.1101/2024.05.31.596898

**Authors:** Eugene F. Baulin, Davyd R. Bohdan, Dawid Kowalski, Milena Serwatka, Julia Świerczyńska, Zuzanna Żyra, Janusz M. Bujnicki

## Abstract

The functions of non-coding RNAs are largely defined by their three-dimensional structures. RNA 3D structure is organized hierarchically and consists of recurrent building blocks called tertiary motifs. The computational problem of RNA tertiary motif search remains largely unsolved, as standard approaches are restrained by sequence, interaction network, or backbone topology. We developed the ARTEM superposition algorithm, which is free from these limitations. Here, we present a version of ARTEM that allows automated searches of RNA and DNA structure databases to identify 3D structure motifs. We exemplify it by a search of motifs isosteric to the kink-turn motif. This widespread motif plays a role in many aspects of RNA function, and its mutations are known to cause several human syndromes. With ARTEM, we discovered two new kink-turn topologies, multiple no-kink variants of the motif, and showed that a ribosomal junction in bacteria forms either a kink or a no-kink variant depending on the species. Additionally, we identified kink-turns in the catalytic core of group II introns, whose structures have not previously been characterized as containing kink-turns. ARTEM opens a fundamentally new way to study RNA and DNA 3D folds and motifs and analyze their correlations and variations.

## MAIN

The functions of non-coding RNAs are largely defined by their three-dimensional structures [1]. RNA 3D structure is organized hierarchically and can be seen as a set of building blocks, RNA 3D modules [2]. The recurrent modules that preserve their features in various RNAs across different structural contexts are called RNA tertiary motifs [3]. Commonly, RNA tertiary motifs are classified into two major classes: local loop motifs, nested within elements of RNA secondary structure [4], such as kink-turns [5], and long-range motifs formed between distant structural segments, such as D-loop/T-loop interaction motifs [6]. Long-range motifs remain much less explored [7, 8] compared to loop motifs [4].

Among the main reasons for the lag in long-range motif exploration are the limitations of existing computational methods designed for the motif search problem, which is formulated as the task of identifying instances of a given 3D motif within a given 3D structure or a set of structures [9-14]. The commonly used methods are limited by at least one of three major restraints: sequence (types of bases) [10, 11], annotations of interactions (e.g., types of base pairs [15]) [9, 11, 12, 14], and backbone topology (types of loops [16]) [10, 12, 13]. Sequence restraints may hide rare non-canonical motif variants, such as the U-minor variant of the A-minor interaction motif [17]. Searching by interaction network relies on pairwise interaction annotations that differ significantly depending on the tool used [8]. The topology restraints pose the greatest limitation, resulting in just a few examples of structural similarities identified to date between motifs of different topologies [8, 17-19]. Although tools for topology-independent nucleic acid 3D structure superposition exist, they are designed for the global 3D structure alignment problem and are not suitable for motif search [20-22]. These limitations cause current approaches to favor loop motifs over long-range motifs and substantially hinder the identification of structural similarities between the motifs formed within different backbone topology contexts.

The kink-turn motif [23], one of the most well-studied RNA tertiary motifs [5, 24], exemplifies the consequences of the aforementioned limitations. Kink-turns are typically characterized by a canonical stem (C-stem) followed by a bulge that forms a sharp kink in the backbone and a non-canonical stem (NC-stem) starting with *trans-Sugar-Hoogsteen* (*tSH*) *G-A* base pairs [5] (Figure 1). The kink-turn architecture of an internal loop was discovered in one of the first experimentally resolved ribosomal RNA 3D structures in 2001 [23]. The kink-turn architecture of a three-way junction loop was discovered only in 2011 [18] and later described as the k-junction motif in 2014 [19], despite its multiple instances in the same 3D structures of ribosomal RNAs [19]. It was shown that kink-turns are extremely widespread, with examples in spliceosomal RNAs [25], group I introns [26], and riboswitches [19], playing roles in many aspects of RNA function, such as serving as a protein binding site [27]. Mutations disrupting the kink-turn structure in the spliceosomal U4atac RNA have been shown to cause various diseases, such as Taybi–Linder syndrome [25] and Roifman syndrome [28]. Although the resemblance between kink-turns and diverse A-minor junctions was briefly mentioned as early as 2011 [18], the variety of discovered kink-turns remains limited to the two mentioned topologies [5].

**Figure 1.**
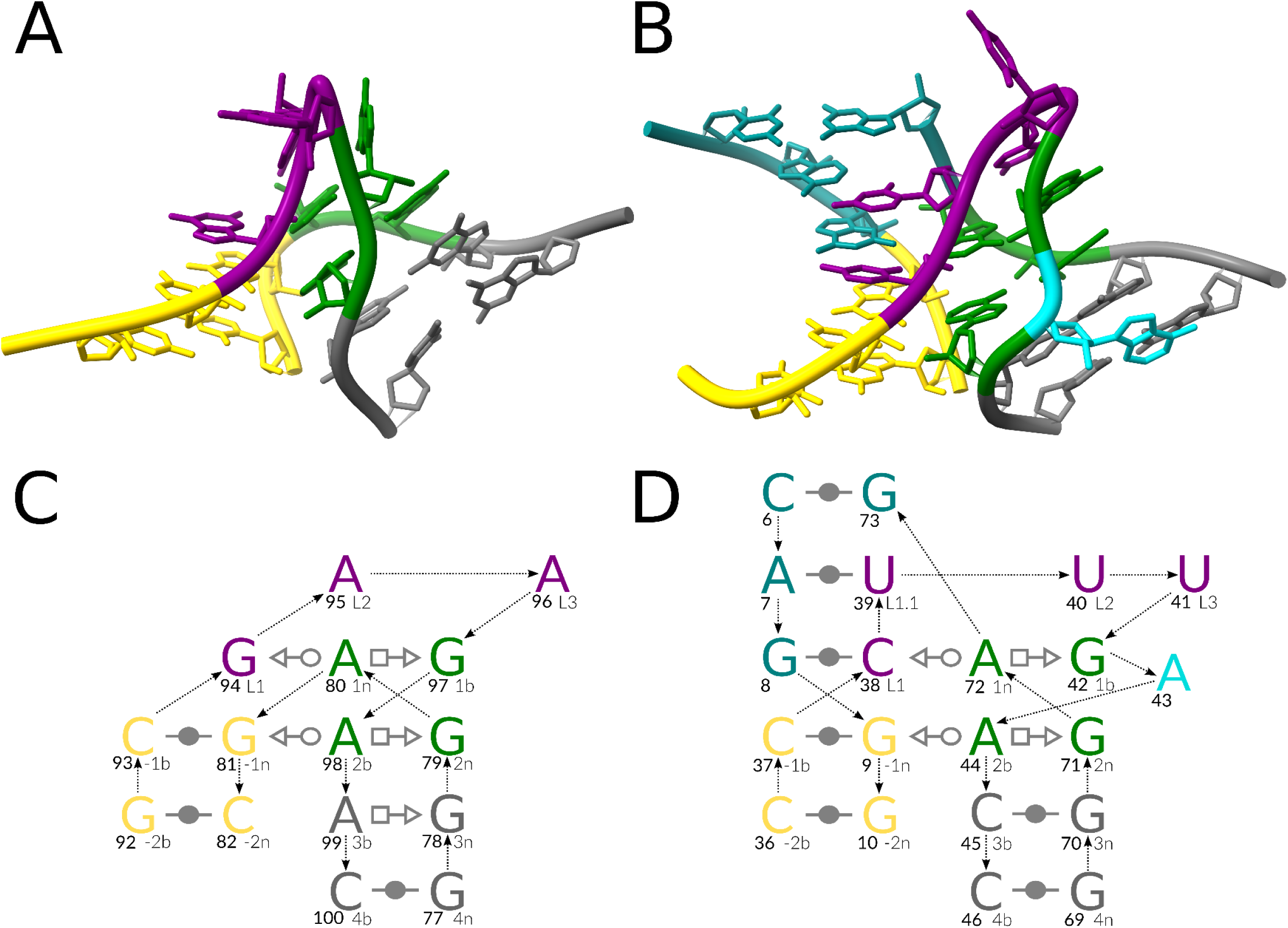
Reference kink-turn motifs. (A) A representative 3D structure of the canonical Kt7 kink-turn [5], LSU rRNA, PDB entry 1FFK, chain 0. (B) A representative 3D structure of the k-junction [5], TPP riboswitch, PDB entry 3D2G, chain A. The interaction schemes of the (C) kink-turn and (D) the k-junction follow the commonly used nomenclature [5]: the canonical stem (C-stem) in gold, the non-canonical stem (NC-stem) in green and gray, the residues of the kink in purple, the third stem (T-stem) and other stems in teal, and the looped-out residues in dark turquoise. The base pair representation follows the Leontis-Westhof (LW) classification [15]. The backbone connectivity is depicted with arrows. Each base is marked with its number in the chain (in bold, left) and its named position in the kink-turn (right). The 3D representations were prepared with ChimeraX (https://www.cgl.ucsf.edu/chimerax/). The 2D representations were prepared with Inkscape (https://inkscape.org/).

Recently, we developed ARTEM [8], a method for sequentially unconstrained superposition of RNA 3D modules. Here, we present a new version of the ARTEM approach adapted to the tertiary motif search problem. We demonstrate the practical utility of the algorithm to identify motifs structurally similar to the kink-turn motif. We identified two new kink-turn topologies: a five-way junction and an external loop. We also discovered several kink-turn-like motifs, including A-minor junctions [18] and long-range motifs [8], that preserve the core interactions of a kink-turn but do not involve the kink. Finally, we demonstrate that the 23S rRNA J94/99 junction, previously identified as a k-junction in *H.marismortui* [19], adopts the no-kink junction topology in other bacteria with a slight distortion of the core kink-turn interactions. Additionally, we identified the coordination loop in group II introns as a kink-turn, although group II intron structures have not been previously characterized as containing kink-turns. Using a benchmark set of group II intron structures, we demonstrated that ARTEM outperforms existing motif search tools. Furthermore, we used ARTEM to search for four types of RNA and DNA tertiary motifs, demonstrating its suitability for structural motifs of varying nucleic acid types, sizes, interaction networks, and backbone connectivities. ARTEM can be uniformly applied to search for both local and long-range motifs, and its advantages open a fundamentally new way to study nucleic acid 3D folds and motifs, and to analyze their correlations and variations.

## RESULTS

We developed ARTEM 2.0 (https://github.com/david-bogdan-r/ARTEM), a new version of the ARTEM algorithm tailored to the nucleic acid tertiary motif search problem. This tool enables automated searches of RNA and DNA structure databases against multiple query modules. ARTEM is equipped with a rich set of user-defined parameters to specify particular 3D structure regions of interest, impose restraints on the identified motifs, and save the superimposed matches or entire nucleic acid-containing input complexes in PDB or mmCIF format. ARTEM advances beyond the limitations of the existing tools by exclusively relying on motif isostericity. It applies to any chosen reference motif, without differentiation between loop motifs and long-range motifs. We exemplified its capabilities by searching for kink-turn-like RNA tertiary motifs. Additionally, we demonstrated ARTEM’s broad applicability by searching for well-known nucleic acid structural motifs, such as G-quadruplex, GNRA-tetraloop, and i-motif, as well as a unique RNA motif with four parallel base pairs recently described [29].

### New backbone topology variants of the kink-turn motif

To search for instances of the kink-turn motif, we selected the representatives of its two known backbone topology variants (Figure 1, Table 1): an instance of the kink-turn internal loop from a 23S rRNA [30] (module #1) and an instance of the k-junction variant from a TPP riboswitch [31] (module #2). We executed ARTEM for the two instances against all RNA-containing entries in the Protein Data Bank [32], see the Methods section for details. The results included 26,564 matches. We annotated the matches with their backbone topology characteristics to facilitate subsequent analysis.

**Table 1.** Representative instances of the kink-turn motif variants.

| # | PDB entry, chain | Description | Topology | RNA molecule | L1 residue | 1n residue | Variant | Figure |
| --- | --- | --- | --- | --- | --- | --- | --- | --- |
| 1 | 1FFK chain 0 | kink-turn Kt7 reference | internal loop | 23S rRNA | G94 | A80 | kink | 1A, 1C |
| 2 | 3D2G chain A | k-junction reference | 3-way junction | TPP riboswitch | C38 | A72 | kink | 1B, 1D |
| 3 | 7P7T chain A | J4/5 | 5-way junction | 23S rRNA | G48 | A117 | kink | 2A, 2B |
| 4 | 7PKT chain 8 | [J99/101] | external loop | L8 rRNA | A453 | A481 | kink | S1A, S1C |
| 5 | 7ZHG chain 2 | no kink but stem | 3-way junction | 16S rRNA | G1073 | A1032 | no-kink | S2A, S2C |
| 6 | 8EUY chain 1 | no kink but two stems | 5-way junction | 28S rRNA | G973 | A685 | no-kink | S2B, S2D |
| 7 | 6ME0 chain A | no kink but stem | 4-way junction | group II intron | C141 | A181 | no-kink | 3A, 3C |
| 8 | 7PWO chain 1 | long-range motif | helix-helix interface | 28S rRNA | C1572 | G2651 | no-kink | 3B, 3D |
| 9 | 4V8O chain BA | J99/101 | 3-way junction | 23S rRNA | G2830 | A2883 | kink | S1B, S1D |
| 10 | 3CC2 chain 0 | J94/99 | complex pseudoknotted junction | 23S rRNA | G2823 | A2914 | kink | 4A, 4C |
| 11 | 4V8O chain BA | J94/99 | complex pseudoknotted junction | 23S rRNA | C2788 | A2892 | no-kink | 4B, 4D |

Via manual inspection of the results, we identified representative instances of two new backbone topology variants of the kink-turn motif (Table 1): a five-way junction in a 23S rRNA [33] (module #3, Figure 2) and an external loop in a mitochondrial LSU rRNA fragment [34] (module #4, Supplementary Figure S1A, C). Below, we use the commonly accepted nomenclature of the kink-turn structure [5], see the Methods section for details.

**Figure 2.**
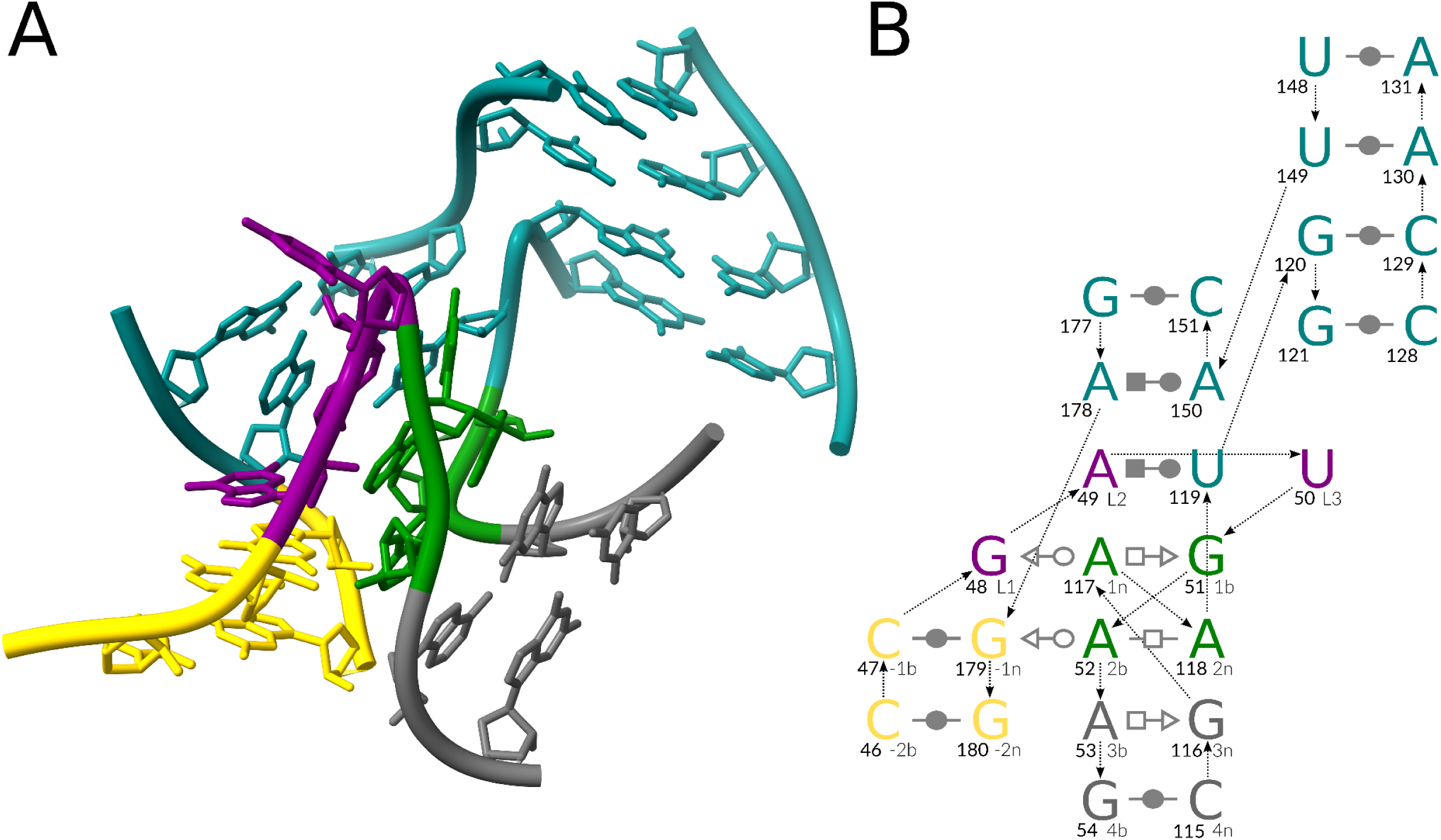
Representative J4/5 five-way k-junction module. (A) A 3D structure of the J4/5 five-way k-junction, LSU rRNA, PDB entry 7P7T, chain A. (B) A 2D interaction scheme of the junction: the canonical stem (C-stem) in gold, the non-canonical stem (NC-stem) in green and gray, the residues of the kink in purple, and the other stems in teal. The base pair representation follows the Leontis-Westhof (LW) classification [15]. The backbone connectivity is depicted with arrows. Each base is marked with its number in the chain (in bold, left) and its named position in the kink-turn (right). The 3D representation was prepared with ChimeraX (https://www.cgl.ucsf.edu/chimerax/). The 2D representation was prepared with Inkscape (https://inkscape.org/).

We mapped the identified five-way junction (module *#3*) to the *J4/5* region of the 23S rRNA structure. The J4/5 region was previously reported to form a k-junction [19], but it was incorrectly identified as a three-way junction. The interaction network of the J4/5 region is closer to the reference module #1, with a *tSH G-A* base pair in position *3n-3b,* but it has a non-linear matching of residues *1n* and *2n,* resulting in a *tHH 2n-2b* base pair.

The external loop (module *#4*) was mapped to the *J99/101* region of the 23S rRNA, which had not been previously reported as a kink-turn or a k-junction. The *J99/101* region forms a canonical three-way k-junction architecture in non-mitochondrial 23S rRNAs (module #9, Supplementary Figure S1B, D). Its interaction network is very close to that of the reference module #2, with one substantial difference: the *2n-2b* base pair is a canonical *cWW* base pair, which is incompatible with the *tSW* interaction formed between residues *-1n* and *2b* in the reference modules.

### No-kink variants of the kink-turn motif

We observed matches of several unique backbone topologies that lacked the kink while exhibiting the C-stem/NC-stem arrangement isosteric to that of the kink variants (modules #5-#8, Table 1, Figure 3, Supplementary Figure S2). Three of these modules (#5-#7) were of different junction loop architectures, all of which can be classified as A-minor junctions, whose resemblance to kink-turns was reported previously [18]. In contrast, module #8 is not a loop but a helix-helix interface, showcasing the long-range kink-turn-like motif for the first time.

**Figure 3.**
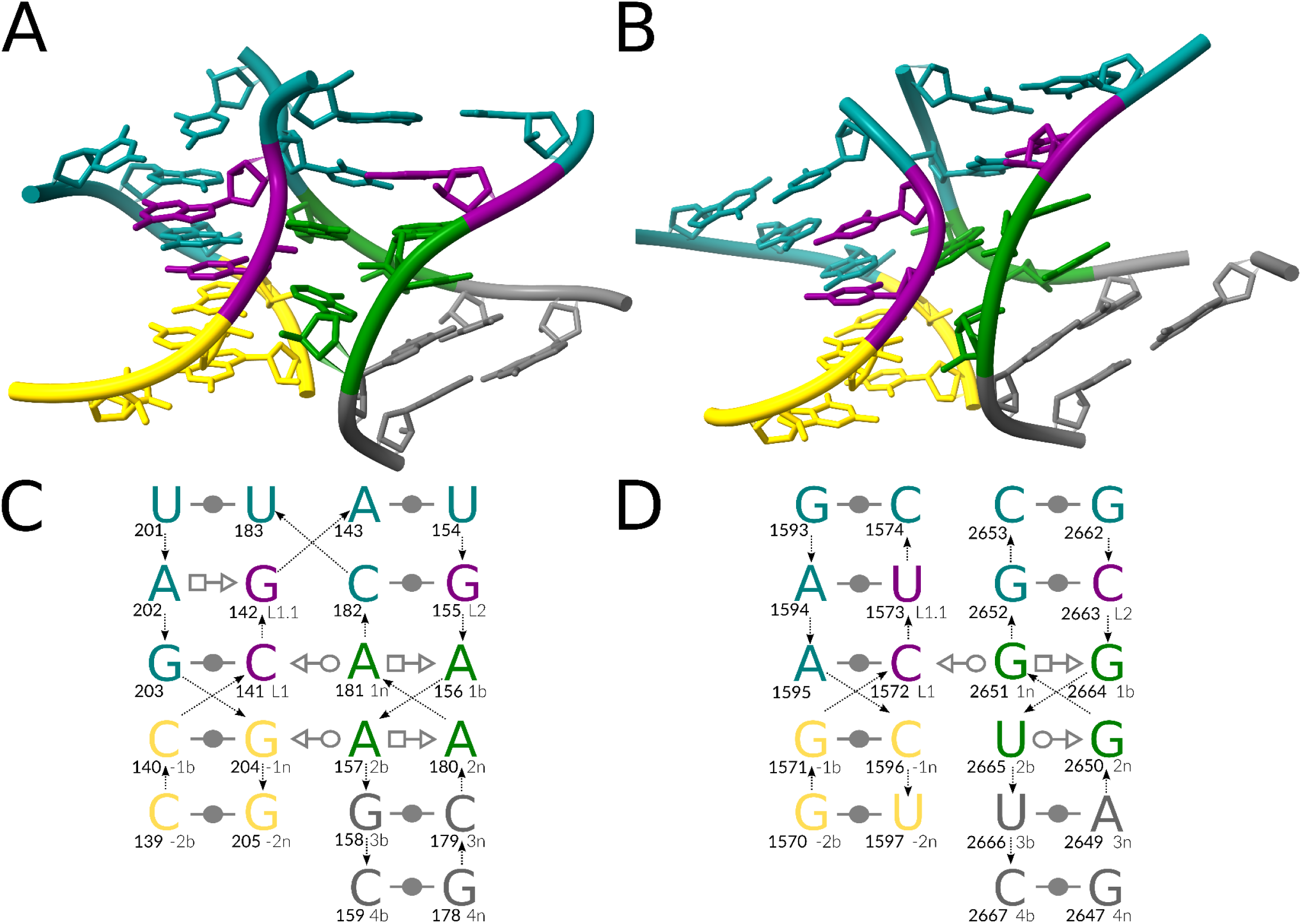
Representative no-kink variants #7 and #8. (A) A 3D structure of the four-way A-minor junction, group II intron, PDB entry 6ME0, chain A. (B) A 3D structure of the long-range helix-helix interface, 28S rRNA, PDB entry 7PWO, chain 1. The interaction schemes of (C) the four-way junction and (D) the long-range motif: the canonical stem (C-stem) in gold, the non-canonical stem (NC-stem) in green and gray, the residues matching the kink in purple, and the other residues in teal. The base pair representation follows the Leontis-Westhof (LW) classification [15]. The backbone connectivity is depicted with arrows. Each base is marked with its number in the chain (in bold, left) and its named position in the kink-turn (right). The 3D representations were prepared with ChimeraX (https://www.cgl.ucsf.edu/chimerax/). The 2D representations were prepared with Inkscape (https://inkscape.org/).

The no-kink three-way junction (module #5, Supplementary Figure S2A, C) and the no-kink five-way junction (module #6, Supplementary Figure S2B, D) both have a pyrimidine in position *1b* instead of the commonly found guanosine. While this is the only notable difference for module #6, module #5 also features a pyrimidine in position *2n* and a non-linear matching of residues *1b* and *2n,* with *1b* belonging to the non-bulge strand.

The no-kink four-way junction (module #7, Figure 3A, C) is the closest module to the reference variants in terms of interactions. The long-range motif (module #8, Figure 3B,D) is the only match that features a non-adenosine (guanosine) *1n* residue. It also includes a uridine in *syn* conformation at position *2b* that does not interact with the *-1n* residue. Notably, there is a striking 3D structure similarity between modules #7 and #8 (Figure 3A,B). Their four RNA strands are arranged similarly but form canonical base pairs with different partners (see the top two base pairs in Figure 3), resulting in distinct topologies: a four-way junction and a long-range helix-helix interface.

### The identified kink/no-kink switch

We discovered that the *J94/99* region of 23S rRNA, another ribosomal k-junction identified previously in *H.marismortui* [19] (module #10, Figure 4A, C), adopts the no-kink junction topology in other bacteria (module #11, Figure 4B, D) with a slight distortion of the core kink-turn interactions (Figure 4D).

**Figure 4.**
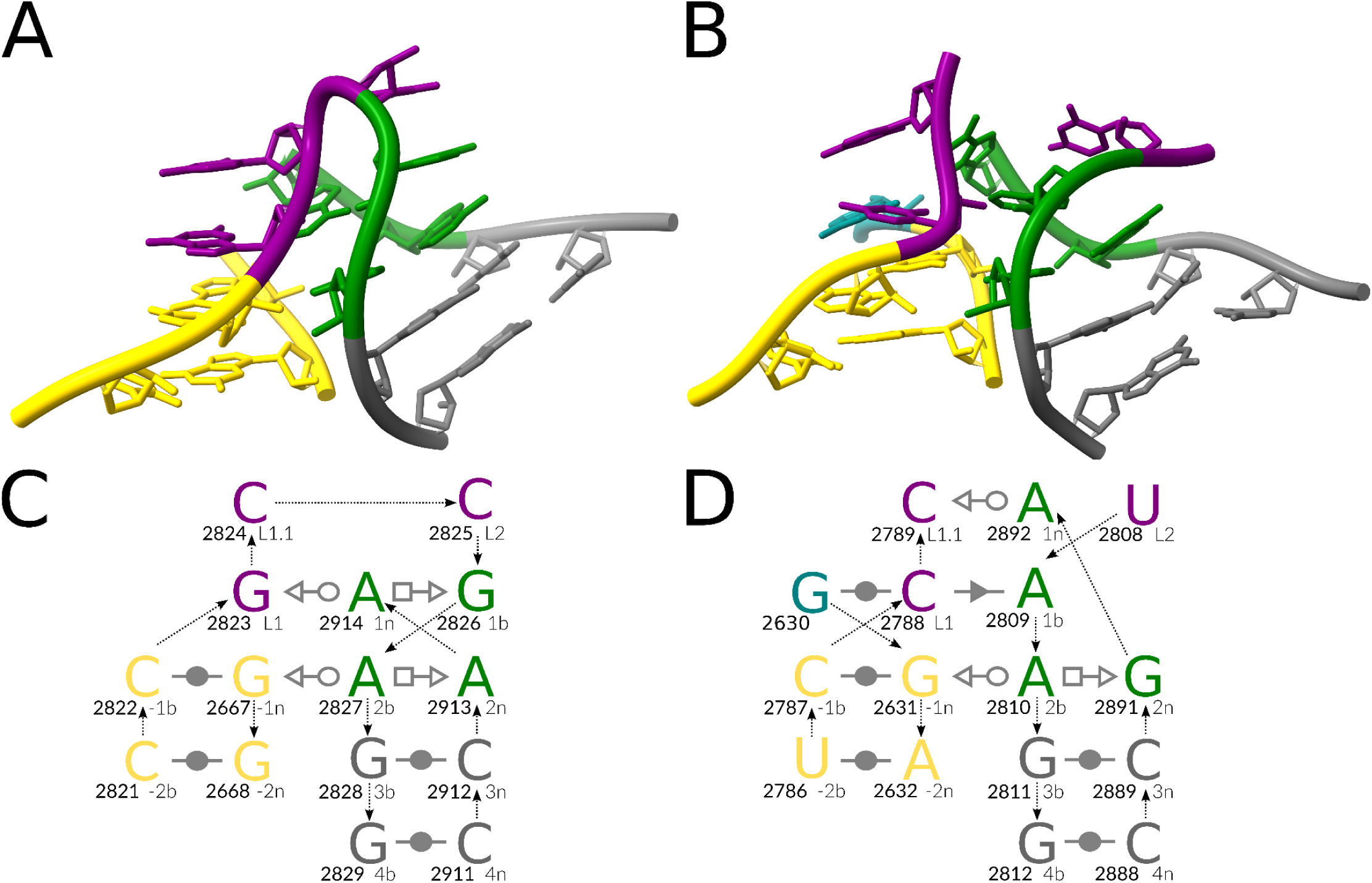
Representative J94/99 region modules. (A) A 3D structure of the J94/99 k-junction in H.marismortui, 23S rRNA, PDB entry 3CC2, chain 0. (B) A 3D structure of the J94/99 no-kink variant, LSU rRNA, PDB entry 4V8O, chain BA. The interaction schemes of (C) the J94/99 k-junction and (D) the no-kink variant: the canonical stem (C-stem) in gold, the non-canonical stem (NC-stem) in green and gray, the residues of the kink in purple, and the other residues in teal. The base pair representation follows the Leontis-Westhof (LW) classification [15]. The backbone connectivity is depicted with arrows. Each base is marked with its number in the chain (in bold, left) and its named position in the kink-turn (right). The 3D representations were prepared with ChimeraX (https://www.cgl.ucsf.edu/chimerax/). The 2D representations were prepared with Inkscape (https://inkscape.org/).

In module #11, Helix 98, absent from the *H.marismortui* 23S rRNA [35], is inserted between residues *L1.1* and *L2* that form the kink in module #10. The *1n* adenosine in the no-kink variant is pushed out of its position by residue *1b* and interacts with residue *L1.1* instead of *L1* (Figure 4D). Also, both modules #10 and #11 formally belong to complex pseudoknotted junction loops, which may be considered yet another backbone topology variant.

### Conservation analysis of ribosomal k-junctions

We analyzed the sequence conservation of the three k-junctions identified in 23S rRNAs (Supplementary Figure S3). The conservation patterns differ substantially between the junction regions but align well with the individual characteristics of the motifs.

The *J4/5* region exhibits the largest number of strictly conserved residues, including the *tHS A-G* base pair in position *3b-3n,* which is also characteristic of the canonical Kt7 kink-turn (module #1). The *1n* residue in the *J94/99* region is the least conserved *1n* residue among the junctions, consistent with its distorted position in the presence of *Helix 98* (module #11). Surprisingly, the *2b-2n cWW U-A* base pair is highly conserved in the *J99/101* region, while the *J99/101* C-stem base pairs are the least conserved among the junctions.

### The coordination loop in group II introns is a kink-turn

While manually inspecting the kink-turn matches obtained with ARTEM, we identified a kink-turn of the canonical internal loop architecture in a 3D structure of group IIC intron [36] (Figure 5A). To the best of our knowledge, occurrences of kink-turns in group II intron structures were never previously reported. To close this gap, we selected a set of seven group II introns from the representative set of RNA structures [37] and surveyed them for kink-turns, both manually and using ARTEM and other existing motif search tools [9-14], see the Methods section for details.

**Figure 5.**
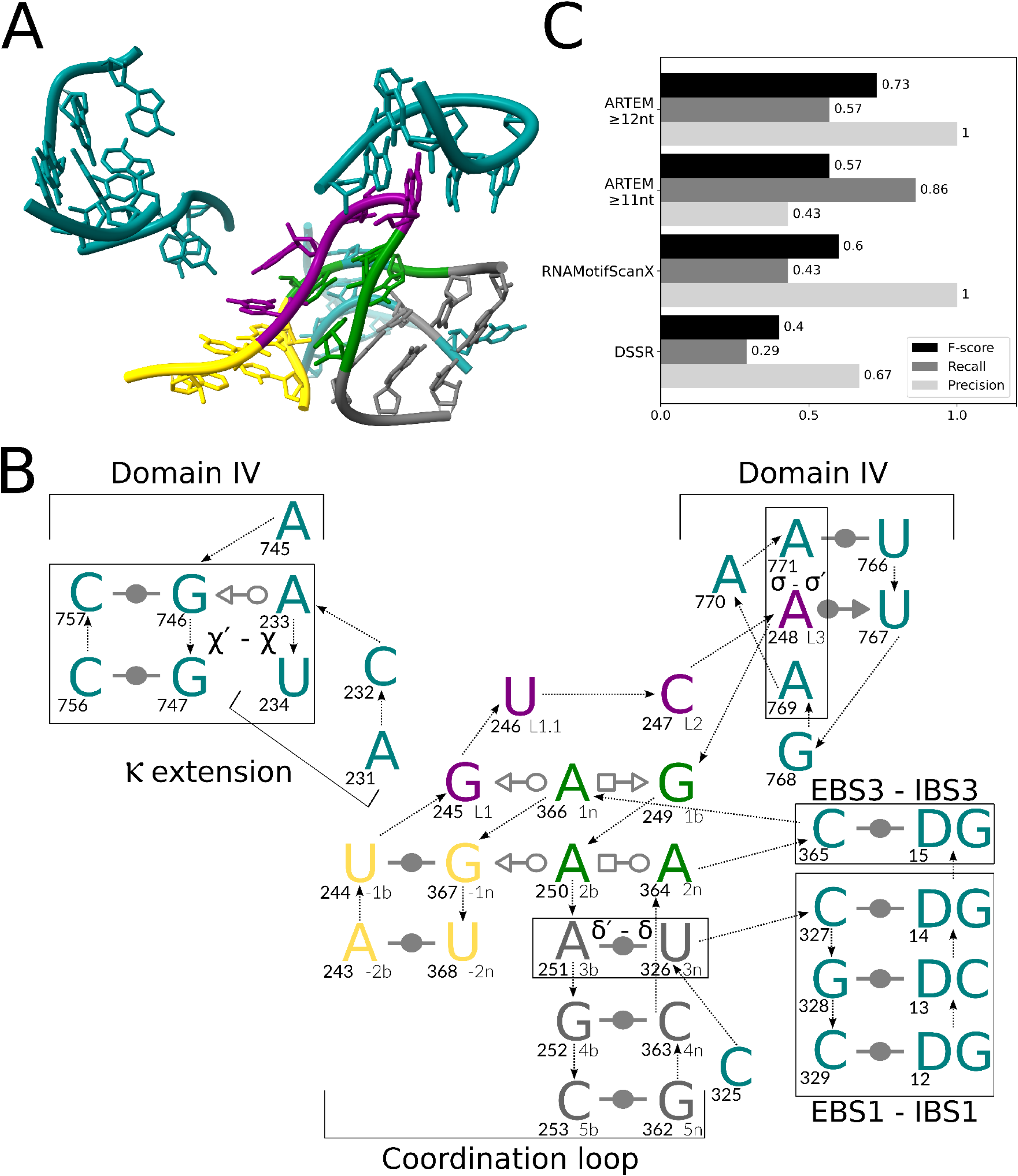
Group IIC intron coordination loop. (A) A representative 3D structure of the coordination loop kink-turn, group IIC intron, PDB entry 6ME0, chain A. (B) The interaction scheme of the kink-turn: the canonical stem (C-stem) in gold, the non-canonical stem (NC-stem) in green and gray, the residues of the kink in purple, and the other residues in teal. The base pair representation follows the Leontis-Westhof (LW) classification [15]. The backbone connectivity is depicted with arrows. Each base is marked with its number in the chain (in bold, left) and its named position in the kink-turn (right). The tertiary interactions (Greek letters) and the intron regions are marked according to the conventional naming [36, 38]. (C) ARTEM benchmarking against other tools that identified at least one of the seven kink-turns in the seven representative group II intron structures. The 3D representation was prepared with ChimeraX (https://www.cgl.ucsf.edu/chimerax/). The 2D representation was prepared with Inkscape (https://inkscape.org/). The barchart was prepared with Matplotlib (https://matplotlib.org/).

We identified seven kink-turn modules in the seven group II intron structures (Supplementary Table S1). Five of the seven kink-turn modules were found in the functional intron region of domain I (DI) known as the coordination loop [38], present in the structures of group IIB and group IIC introns. As implied by its name, the loop coordinates the formation of the intron’s catalytic core by interacting with the Ⲕ extension bulge and the exon binding site 1 (EBS1) loop [38]. Additionally, the coordination loop encompasses the single-residue exon binding site 3 (EBS3). Subsequently, exon-intron recognition is facilitated through interactions with the intron binding sites IBS1 and IBS3 (Figure 5B). The remaining two kink-turns were identified as instances of a single Domain III (DIII) kink-turn module in group IIC introns.

The kink-turn architecture is crucial for the functioning of the coordination loop (Supplementary Table S1). The residues *L1, L1.1*, and *L2* interact with the Ⲕ extension. The *L3* residue forms a base wedged element (BWE) [39] with a Domain IV (*DIV*) loop prior to DNA integration [36]. In the group IIC intron structures, the residues *L2, L3, 1b*, and *EBS3* are exposed to the reverse transcriptase protein. Furthermore, the unusual long-range δ′-δ base pair in position *3b-3n* facilitates the stacking between *EBS3* and *EBS1*. Notably, the two representative group *IIA* intron structures lack *EBS3*, and their coordination loops do not form a kink-turn.

Surprisingly, the coordination loop was never identified as a kink-turn previously. Possibly, the long-range δ′-δ base pair replacing the local *3b-3n* base pair and the looped-out *EBS3* residue hid the motif from identification via sequence- and backbone-restrained kink-turn searches. Among the previously existing motif search tools, only DSSR and RNAMotifScanX were able to identify at least one kink-turn in the seven selected 3D structures. DSSR, performing the sequence-based search, identified only the two DIII kink-turn modules. RNAMotifScanX, searching with the consensus kink-turn interaction network, identified the two DIII kink-turns and a single coordination loop kink-turn. In contrast, ARTEM, searching with a single Kt7 reference module, identified four of the five coordination loop kink-turns with the 12nt threshold on the match size. With the 11nt threshold, ARTEM identified all five coordination loop kink-turns and one of the DIII kink-turns, along with eight false positive matches. Consequently, ARTEM outperformed the existing tools, demonstrating an impressive recall of 57% at 100% precision with the 12nt threshold, and 86% recall at 43% precision with the 11nt threshold (Figure 5C).

### ARTEM can detect various types of RNA and DNA tertiary motifs

To demonstrate the broad applicability of ARTEM, we also used it to search for four types of nucleic acid tertiary motifs across all RNA- and DNA-containing PDB entries. This search against complete but redundant datasets aimed to verify the characteristic features of the motifs and to define optimal sequence and RMSD restraints for ARTEM use. Three of the four motifs are well-known: the parallel G-tetrad [40] (Supplementary Figure S4A), GNRA-tetraloop [41] (Supplementary Figure S5), and i-motif [42] (Supplementary Figure S6). The fourth is an unusual motif of four base pairs formed between parallel-oriented strands (Supplementary Figure S7), recently identified in place of a predicted pseudoknot in the crystal structure of cap-independent translation enhancers (CITE) from Pea enation mosaic virus RNA 2 (PEMV2) [29]. We refer to this motif as the parallel-pairing motif.

Distinct RMSD distributions of all-guanine and non-all-guanine matches (Supplementary Figure S4B, C) confirmed the high specificity of the parallel G-tetrad motif to guanosine bases, with only a few non-guanine matches identified with RMSD < 1.0 Å, such as an all-adenine tetrad (Supplementary Figure 4D). In contrast, at RMSD from 1.0 to ∼1.75 Å, we observed a notable portion of false all-guanine matches that spanned residues from several adjacent non-parallel tetrads (Supplementary Figure 4E, F). Therefore, an RMSD threshold of 1.0 Å is recommended when searching for parallel tetrads with ARTEM, with additional sequence restraints if necessary.

Unexpectedly, among GNRA-tetraloop matches identified at RMSD < 1.0 Å, we did not observe clear separation between RMSD distributions of GNRA and non-GNRA matches (Supplementary Figure S5B). Among these matches, 6.8% were GAAG tetraloops, and 4.7% were UAAC tetraloops, which adopted the same fold but did not conform to the GNRA pattern. To account for redundancy, we analyzed non-coding RNA families containing these matches. The GAAG loop was found exclusively in ribosomal RNAs (both SSU and LSU, across seven RNA families) and was further stabilized either by RNA-protein interactions (LSU) or by a base-phosphate interaction (SSU, see Supplementary Figure S5D). The UAAC loop was identified only in a lariat capping ribozyme and in bacterial SSU rRNA, where it was stabilized by a distant adenosine base in both cases (Supplementary Figure S5E). As expected, non-GNRA loops require additional stabilization to form a GNRA-like fold. Interestingly, ARTEM identified a single GNRA-tetraloop match formed by a DNA chain, indicating that this motif is not exclusive to RNA (Supplementary Figure S5C). Overall, an RMSD threshold of 1.5 Å is recommended when searching for GNRA-tetraloops with ARTEM, as matches with distorted folds appear above this threshold (Supplementary Figure S5F).

Among RNA-containing PDB entries, ARTEM identified only 12 matches for the reference RNA i-motif instance at RMSD < 2.0 Å, all being all-cytidine matches at RMSD < 0.35 Å from the same PDB entry as the reference (Supplementary Figure S6C). Thus, PDB entry 1I9K is the only entry containing an RNA i-motif, confirming that i-motifs are generally unfavorable in RNA structures. Among DNA-containing PDB entries, ARTEM identified 144 all-cytidine i-motif matches at RMSD < 1.0 Å, one match involving thymine-thymine base pairs at RMSD = 1.004 Å, and four matches involving adenosine-adenosine base pairs at RMSD = 1.24 Å (Supplementary Figure S6B, D, E).

Across RNA- and DNA-containing PDB entries, ARTEM did not identify a single match for the parallel-pairing motif at RMSD < 1.0 Å. The closest match was found at RMSD = 1.679 Å and did not exhibit the parallel-pairing pattern characteristic of the reference motif (Supplementary Figure S7D, E, F). Therefore, this search confirms the uniqueness of the parallel-pairing motif described in the CITE RNA from PEMV2. These analyses demonstrate ARTEM’s unique suitability for identifying similarities between nucleic acid structural motifs, regardless of the nucleic acid type, size, interaction network, or backbone connectivity.

## DISCUSSION

In this work, we developed ARTEM 2.0, an adaptation of the ARTEM algorithm suitable for tackling the long-standing problem of RNA tertiary motif search. ARTEM relies exclusively on motif isostericity and enables the identification of topology-independent structural similarities between the motifs. Therefore, ARTEM overcomes the “*clear shortcoming in classifying RNA motifs simply based on their secondary structures*” [18]. This method applies to any chosen RNA or DNA 3D module, without differentiation between loop motifs and long-range motifs.

We employed ARTEM to explore the landscape of modules isosteric to the kink-turn motif. We identified several new topological kink-turn variants, including a five-way junction and an external loop. We showed that certain motifs preserve the same core interactions of the kink-turn but lack the kink. We demonstrated that the ribosomal region *J94/99* can form either a kink or a no-kink variant of the motif depending on the species, albeit with a distortion in residue arrangement in the no-kink variant. Furthermore, we identified kink-turns with a complex network of long-range base pairs in the catalytic core of group II introns, whose structures were never previously reported to involve kink-turns. Analyzing the identified representative kink-turn modules, we found that their only consensus feature is the *1n* purine (almost always adenosine) forming the type I A-minor interaction [43] with the residue *L1* (and its partner *L1p* if present) and donating its Hoogsten edge for base-pairing with the residue *1b*. Strikingly, a 3D module can even lack this essential feature and still be largely isosteric to kink-turns, as evidenced by module #11 (Figure 4B,D).

Additionally, we used ARTEM to search for four distinct types of nucleic acid structural motifs: parallel G-tetrad, GNRA-tetraloop, i-motif, and parallel-pairing motif. For example, we demonstrated that GNRA-like folds can be formed by non-GNRA loops with additional stabilization provided by distant residues or other molecules (Supplementary Figure S5). Thus, we showed that ARTEM is readily applicable to structural motifs of various sizes, interaction networks, and backbone connectivities.

A notable limitation of ARTEM is that it may require some fine-tuning based on the particular motif of interest and the desired *FP/FN* ratio. In turn, ARTEM offers high flexibility and enables a rich set of user-defined restraints. Furthermore, ARTEM outperformed existing tools in the search for kink-turns among group II introns, even without the introduction of additional restraints on matches beyond a single match-size threshold (Figure 5C). Nevertheless, we intend to develop a separate tool, tailored for the annotation of the most prevalent RNA and DNA tertiary motifs, that can be used by non-experts as a closed user-friendly solution.

ARTEM introduces a fundamentally new approach to investigating nucleic acid 3D folds and motifs and analyzing their correlations and variations. It extends the concept of isostericity from base pairs [44] to more complex tertiary motifs. By employing a purely isosteric-based tertiary motif search, ARTEM promises to unveil numerous hidden similarities among recurrent RNA and DNA 3D modules, significantly advancing our understanding of the principles governing RNA and DNA 3D structure organization.

## METHODS

### Definitions

Below we provide definitions as applied to RNA 3D structures; however, these definitions apply equally to DNA 3D structures. The definitions of RNA secondary structure elements used in this work follow our previous study [8]. A *stem* is a helical region formed by at least two consecutive stacked base pairs, of Watson-Crick G-C/A-U or wobble G-U type, and a *loop* is a set of single-stranded regions confined by a stem. A loop and a stem are called *adjacent* if at least one residue of the loop is a direct neighbor in sequence of a residue of the stem. Two elements (stems or loops) are called *distant* if they are not adjacent and don’t share common adjacent elements. An *internal loop* includes two strands and is adjacent to two stems. An *N-way junction loop* involves *N > 2* strands and is adjacent to *N* stems. A loop of any type is called *external* if it includes dangling ends among its strands.

An *RNA 3D module* is loosely defined as a set of interacting residues that can be represented with a connected graph. A module is called *long-range* if it involves at least one pair of residues that belong to distant RNA secondary structure elements. Otherwise, a module is called *local*. An *RNA tertiary motif* is defined as a recurrent RNA 3D module. A module is called a *motif instance* if it’s identified as the motif representative. We defined the computational problem of RNA tertiary motif search as the task of identifying instances of a given motif in a query RNA 3D structure having an instance of the motif as the reference.

In this work, we adhere to the established nomenclature of the kink-turn structure [5] and Leontis-Westhof base pair classification [15] (Figure 1). A residue forms a base pair with another residue via one of its three edges: *Watson-Crick* (*W*, circle), *Hoogsteen* (*H*, square), or *Sugar* (*S*, triangle), in either *cis* (filled) or *trans* (empty) orientation. A canonical kink-turn comprises two helical regions, a canonical stem (C-stem, in gold) and a non-canonical stem (NC-stem, in green and gray), connected by an internal loop. The longer strand of the loop (bulge, in purple) involves the sharp kink in its backbone. The NC-stem starts with two characteristic *trans-Sugar-Hoogsteen* (*tSH*) *G-A* base pairs (in green). The residues of the bulge are labeled as *L1*, *L2*, and *L3*. The base pairs of NC-stem are labeled with positive numbers starting from the base pair adjacent to the bulge, with lowercase *b* (bulge side) or *n* (non-bulge side) letters: *1n/1b* base pair, *2n/2b* base pair, and so on. The C-stem base pairs are labeled with negative numbers: *-1b/-1n* base pair, *-2b/-2n*, and so on. The k-junction variant (Figure 1B, D) involves an additional *L1.1* bulge residue and the third stem (T-stem, in teal). The *L1* residue of the k-junction forms a canonical base pair with the residue that we labeled *L1p*.

We should note that the NC-stem, as defined in [5], deviates from the strict stem definition, with the *tSH G-A* base pairs formally falling within the internal loop. To maintain consistency, we adhere to the conventional NC-stem definition, while employing the strict stem definition for RNA secondary structure annotation.

### ARTEM tool improvement

As the original ARTEM tool [8] reports all local structural similarities identified between the two nucleic acid 3D modules, we modified it to make it more suitable for the nucleic acid tertiary motif search. ARTEM 2.0 introduces three new features compared to its predecessor (https://github.com/david-bogdan-r/ARTEM). First, the input structures (both reference and query) can be specified with a folder or mask instead of a single file, allowing one to search against a set of RNA/DNA structures with one command. Second, we added a *nosub* option to remove sub-matches (matches involved in larger matches), allowing one to filter out partial hits. Third, we introduced an *rst* option, enabling users to specify additional restraints of four different types: (i) filtering out matches missing specified residues, (ii) requiring the specified residues to match with particular base types (allowing the IUPAC nomenclature: https://en.wikipedia.org/wiki/Nucleic_acid_notation), (iii) requiring the specified residues to match under a given RMSD threshold, and (iv) requiring the specified residues to match with a continuous nucleic acid strand. The continuity of a strand is verified by O3′-P atom distances under 2.0 *Å* between neighboring residues.

### Kink-turn motif search with ARTEM

To conduct the kink-turn motif search with ARTEM, we selected one representative for each of the two known backbone topology variants (Figure 1, Table 1): the canonical kink-turn internal loop (LSU rRNA, PDB entry 1FFK [30], chain 0, *L1* residue *G94*) and the k-junction variant (TPP riboswitch, PDB entry 3D2G [31], chain A, *L1* residue *C38*). For the search, we confined the motif to the following 14 residues: *L1, L1p, -1b, -1n, -2b, -2n, 1b, 1n, 2b, 2n, 3b, 3n, 4b, 4n*. Other labeled residues were excluded due to their dynamic positioning between the kink-turn instances.

To determine the thresholds for the search, we executed ARTEM using the two reference instances as input. ARTEM identified a correct (consistent with labels) match of 13 residues (*L1p* is absent in the canonical kink-turn) at *RMSD* of *1.837 Å* between the two instances. Accordingly, we set the search for the matches of at least 12 residues (*sizemin = 12*) at RMSD under 2.0 Å (*rmsdmax = 2.0*). Additionally, we imposed a restraint requiring the matches to include the counterparts of all of the following residues: *L1, -1b, -1n, 1b, 1n, 2b, 2n*, as they are deemed most essential for the kink-turn motif.

We then executed ARTEM for each of the two references against all RNA-containing structures in the Protein Data Bank [32] (7,292 PDB entries as of December 7th, 2023). The search concluded within 24 hours on an AMD Ryzen 9 5950X machine equipped with 32 CPU cores and 128 GB RAM. ARTEM identified 11,025 matches for the kink-turn and 18,727 matches for the k-junction. Combining the matches identified against both references resulted in 26,564 unique matches (Supplementary Table S2). Finally, to facilitate subsequent analysis, we used the *urslib2* Python library [8] to annotate RNA secondary structure elements and additional characteristics (e.g., local/long-range, kink/no-kink) of the hits’ residues based on the base pair annotations obtained with DSSR [11] (version 2.0). The kink/no-kink distinction was approximated by the distance in RNA sequence between the *2b* and *L1* residues: the kink label was assigned to the matches with a distance under ten residues.

Additionally, we ran a similar ARTEM search against all DNA-containing PDB entries (10,505 entries as of December 7th, 2023). We confirmed that no kink-turn or k-junction matches were identified within DNA molecules.

### Analysis of the kink-turn motif search results

We manually inspected the motif matches identified with ARTEM, targeting novel backbone topologies, as annotated with *urslib2*. The analysis was performed using ChimeraX [45]. The boundaries of the identified representative modules were defined manually, and the modules were then saved as separate coordinate files (https://github.com/febos/ARTEM-KT).

The naming of the ribosomal k-junctions *J4/5* and *J94/99* followed the original k-junction paper [19]. The name for the *J99/101* k-junction was derived analogously using the RibosomeGallery [35] (http://apollo.chemistry.gatech.edu/RibosomeGallery/). The mapping of the identified motifs to the named ribosomal regions was done based on the original structure papers [33, 34, 46]. We should note that modules #9 (PDB entry 4V8O, *J99/101*) and #10 (PDB entry 3CC2, *J94/99*) were missing from the original list of ARTEM matches (Supplementary Table S2). They could be identified with ARTEM only with relaxed thresholds: 11-residue hit at *RMSD = 1.975 Å* against module #2 and 12-residue hit at *RMSD = 1.958 Å* missing *-1n* counterpart against module #1, respectively.

For the conservation analysis, we mapped the three k-junctions to the Rfam seed alignment of the 23S rRNA family RF02541 [47] (https://github.com/febos/ARTEM-KT/tree/main/sto_logo). The relevant sequence regions were then excised from the alignment and used as input for the WebLogo application [48] (https://weblogo.berkeley.edu/) to construct the motif logos.

### Group II intron survey for kink-turns

For the benchmarking of the motif search tools, we selected all representative RNA structures assigned with the “group II” keyword in the representative set [37] (version 3.312 with the resolution cutoff of 4.0 Å, http://rna.bgsu.edu/rna3dhub/nrlist/release/3.312). We excluded the 70-nt structure (PDB entry 1KXK) as it is a short intron fragment. The remaining set included seven group II intron structures (Supplementary Table S1). We then manually identified seven kink-turn 3D modules in the seven structures: five coordination loops and two DIII kink-turns, all of them with the internal loop architecture.

Overall, we utilized seven different RNA tertiary motif search tools (Supplementary Table S3). Although direct comparisons between the tools are often unfeasible, we approached the analysis from the final user perspective, where we have a reference motif instance and a query RNA structure, aiming to identify motif instances within the structure. The 13 residues (*L1, -1b, -1n, -2b, -2n, 1b, 1n, 2b, 2n, 3b, 3n, 4b, 4n*) of the Kt7 kink-turn module (PDB entry *1FFK*, chain *0*, *L1* residue *G94*) were used as the reference motif instance. For all the tools, we considered the identified match to be a true positive (*TP*) if it overlapped with the correct kink-turn by at least 50% of its residues; otherwise, it was classified as a false positive (*FP*). A correct kink-turn instance not identified by the tool was labeled as a false negative (*FN*). Different matches of the same module were merged together before tallying the *TP, FP*, and *FN* values. The quality metrics were subsequently calculated as follows: precision = *TP / (TP + FP)*, *recall = TP / (TP + FN)*, and *F-score = 2*TP / (2*TP + FP + FN)*.

We ran ARTEM with no RMSD threshold but with two different minimum match-size thresholds: 12 residues or 11 residues. For DSSR [11] (version 2.0, http://forum.x3dna.org/rna-structures/) we derived the kink-turn annotations with the “*Normal kink-turn*” label. For RNAMotifScanX [14] (http://genome.ucf.edu/RNAMotifScanX/RNAMotifScanX-release_v0.0.5_x86-64_rhel.tar.gz) we utilized the built-in kink-turn consensus structure (*models/k-turn_consensus.struct*). WebFR3D [9] (http://rna.bgsu.edu/webfr3d/fr3d.php) was initially executed as a purely geometric search for the reference module, and then with *cWW* symbolic restraints added. JAR3D [10] (http://rna.bgsu.edu/jar3d/) was executed with the sequence input matching the reference kink-turn (*GGGAG*CGAAGAAC*), as it does not accept 3D modules. LocalSTAR3D [12] (http://genome.ucf.edu/LocalSTAR3D/) and CircularSTAR3D [13] (https://github.com/ucfcbb/CircularSTAR3D) were run with the compatible DSSR version 1.5.3. LocalSTAR3D reported multiple 8-residue single-stem matches not involved in the correct kink-turns, which we chose to ignore and not to assign as false positives. Consequently, four tools reported zero matches overall: WebFR3D, JAR3D, LocalSTAR3D, and CircularSTAR3D.

### Analysis of other types of RNA and DNA tertiary motifs

Reference instances of the parallel G-tetrad, GNRA-tetraloop, i-motif, and parallel-pairing motif were manually selected and searched against all RNA- and DNA-containing PDB entries (7,292 and 10,505 entries, respectively, as of December 7th, 2023) using ARTEM (see https://github.com/febos/ARTEM-KT/blob/main/REPRODUCE.md). The following thresholds were set for these searches: for the parallel G-tetrad, a minimum match size of 4 residues and a maximum RMSD of 2.0 Å; for the GNRA-tetraloop, a minimum match size of 6 residues (tetraloop plus flanking base pair) with the match required to form a single continuous strand; for the i-motif, a minimum match size of 8 residues; and for the parallel-pairing motif, a minimum match size of 8 residues.

## DATA AVAILABILITY

ARTEM 2.0 is available at https://github.com/david-bogdan-r/ARTEM (DOI: 10.5281/zenodo.14034347). The data on kink-turn modules are available at https://github.com/febos/ARTEM-KT (DOI: 10.5281/zenodo.14034295).

## ACKNOWLEDGMENTS

The authors acknowledge the GIFTED Programme of the Polish Children’s Fund (https://fundusz.org/en/gifted-programme/) for facilitating the visit of talented students to IIMCB in Warsaw, which contributed to this work.

## SUPPLEMENTARY MATERIALS

**Supplementary Figure S1.**
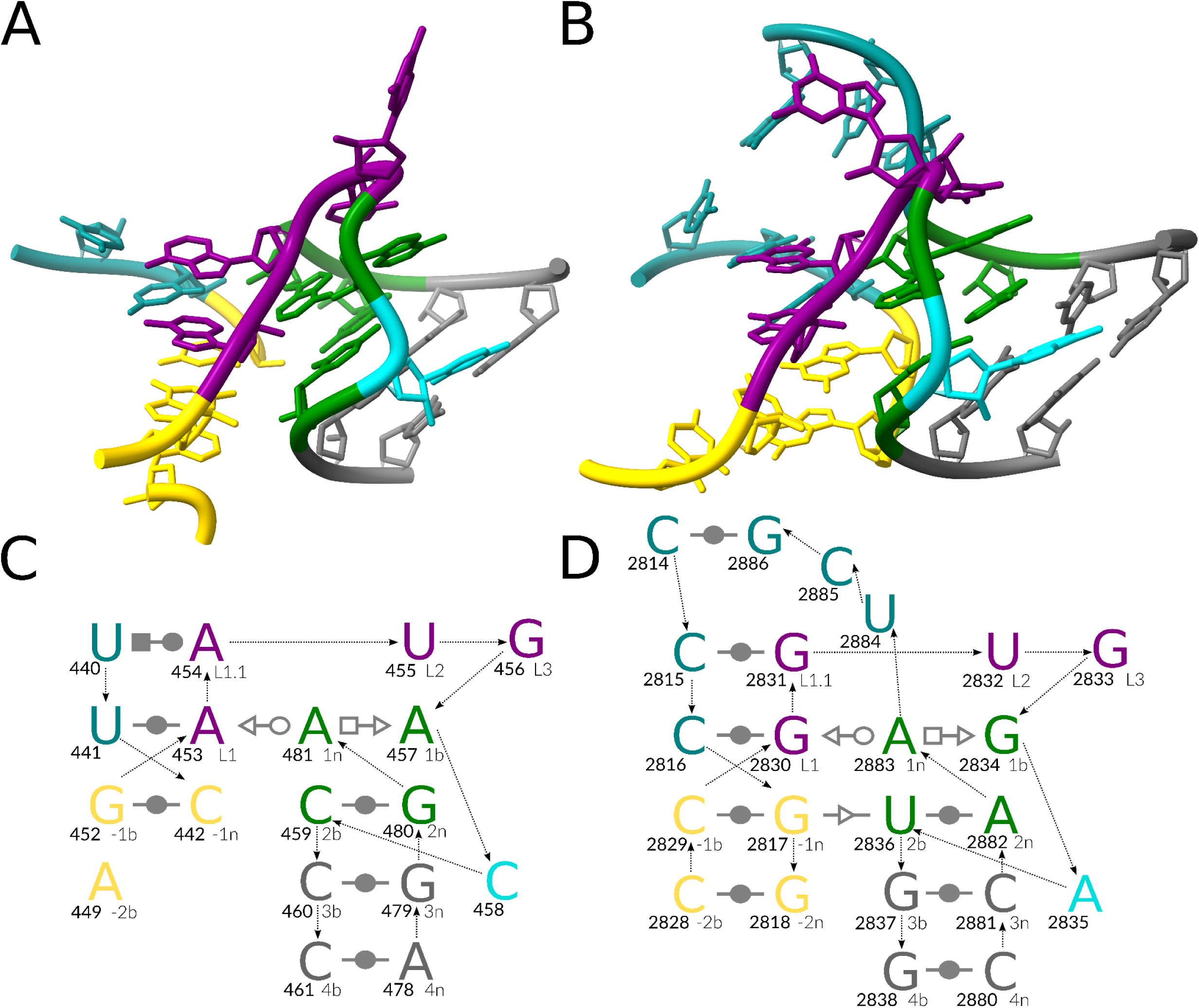
Representative J99/101 k-junction modules. (A) A 3D structure of the J99/101 k-junction with an external loop architecture, L8 rRNA fragment, PDB entry 7PKT, chain 8. (B) A 3D structure of the J99/101 k-junction with a three-way junction architecture, LSU rRNA, PDB entry 4V8O, chain BA. The interaction schemes of (C) the external loop and (D) the three-way junction: the canonical stem (C-stem) in gold, the non-canonical stem (NC-stem) in green and gray, the residues of the kink in purple, the looped-out residues in dark turquoise, and the other residues in teal. The base pair representation follows the Leontis-Westhof (LW) classification [15]. The backbone connectivity is depicted with arrows. Each base is marked with its number in the chain (in bold, left) and its named position in the kink-turn (right). The 3D representations were prepared with ChimeraX (https://www.cgl.ucsf.edu/chimerax/). The 2D representations were prepared with Inkscape (https://inkscape.org/).

**Supplementary Figure S2.**
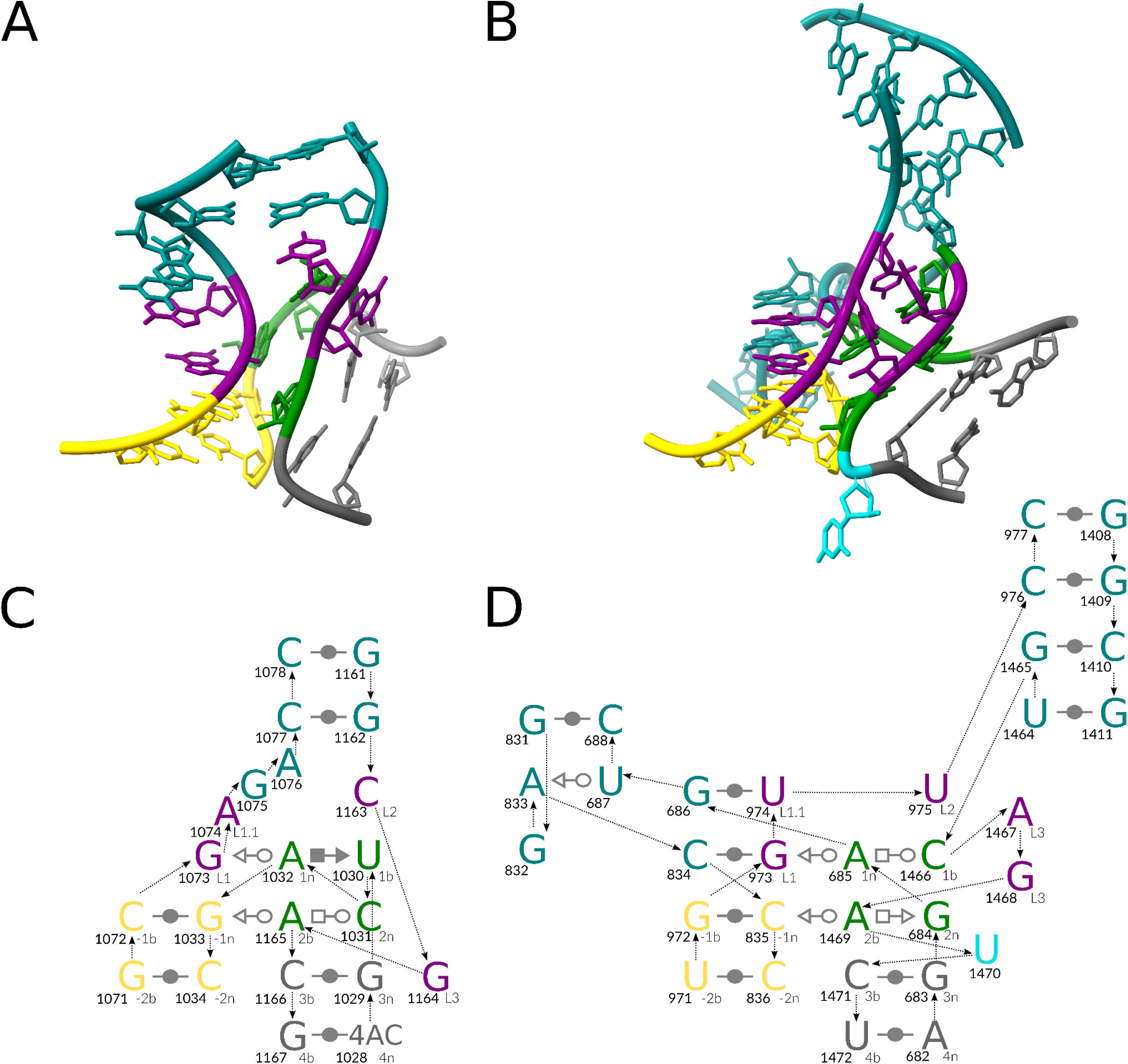
Representative no-kink variants #5 and #6. (A) A 3D structure of the three-way A-minor junction, 16S rRNA, PDB entry 7ZHG, chain 2. (B) A 3D structure of the five-way A-minor junction, 28S rRNA, PDB entry 8EUY, chain 1. The interaction schemes of (C) the three-way junction and (D) the five-way junction: the canonical stem (C-stem) in gold, the non-canonical stem (NC-stem) in green and gray, the residues matching the kink in purple, the looped-out residues in dark turquoise, and the other residues in teal. The base pair representation follows the Leontis-Westhof (LW) classification [15]. The backbone connectivity is depicted with arrows. Each base is marked with its number in the chain (in bold, left) and its named position in the kink-turn (right). The 3D representations were prepared with ChimeraX (https://www.cgl.ucsf.edu/chimerax/). The 2D representations were prepared with Inkscape (https://inkscape.org/).

**Supplementary Figure S3.**
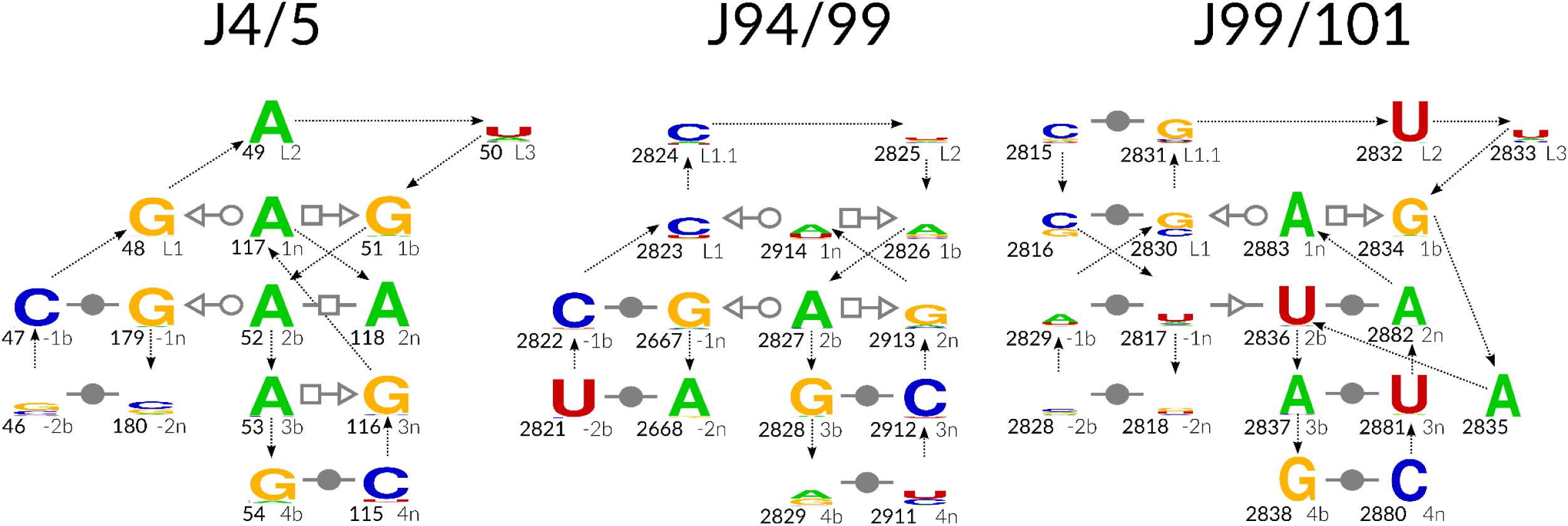
Motif logos of the 23S rRNA k-junctions. The J4/5 junction logo follows the scheme of module #3, PDB entry 7P7T, chain A. The J94/99 junction logo follows the scheme of module #10, PDB entry 3CC2, chain 0. The J99/101 junction logo follows the scheme of module #9, PDB entry 4V8O, chain BA. The base pair representation follows the Leontis-Westhof (LW) classification [15]. The backbone connectivity is depicted with arrows. Each base is marked with its number in the chain (in bold, left) and its named position in the kink-turn (right). The 2D representations were prepared with Inkscape (https://inkscape.org/) and WebLogo (https://weblogo.berkeley.edu/).

**Supplementary Figure S4.**
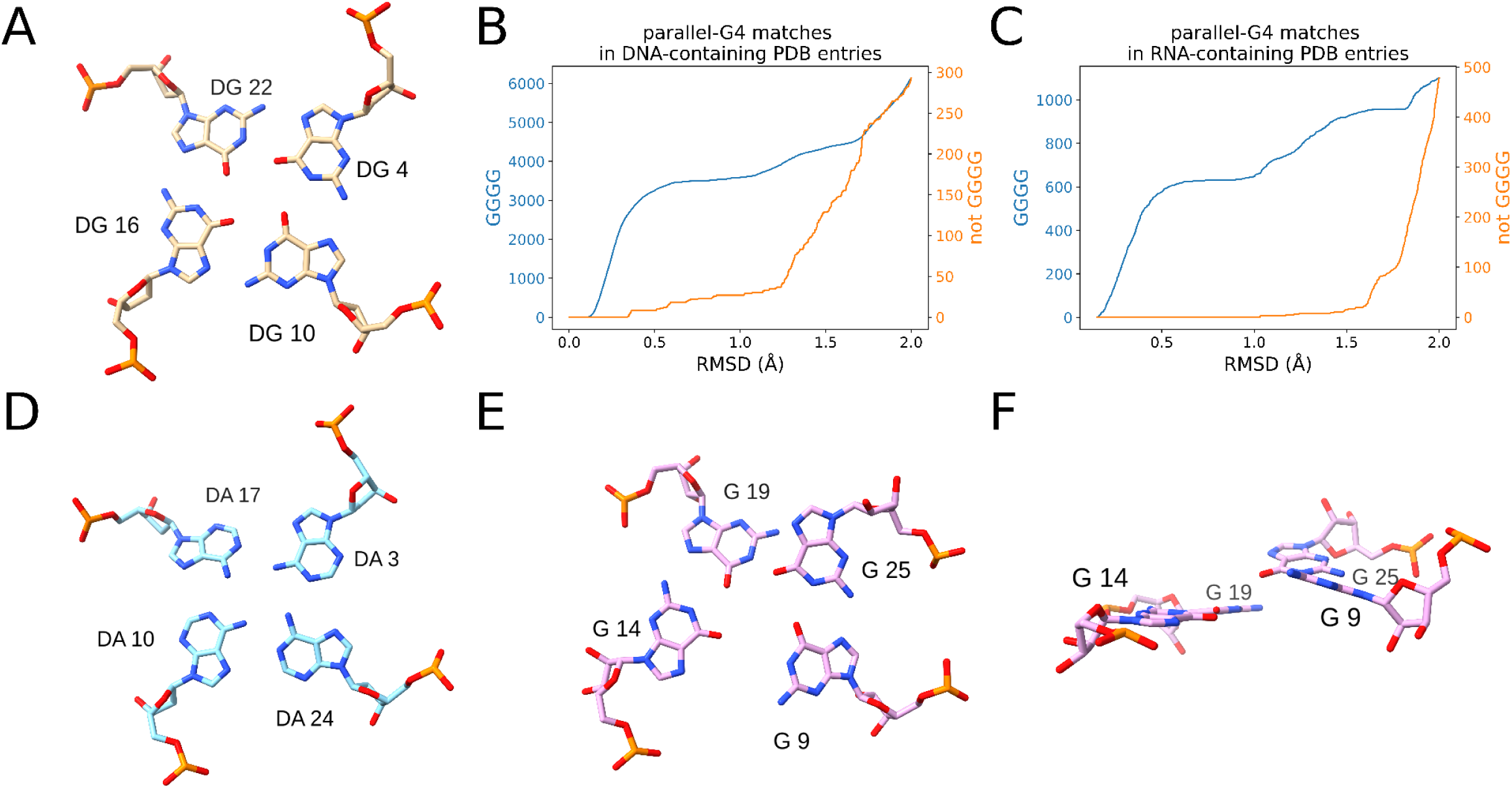
ARTEM search for the parallel G-tetrad motif. (A) Reference parallel G-tetrad instance from a telomeric G-quadruplex, PDB entry 8D79, chain A. (B) all-G (in blue) and non-all-G (in orange) matches of the motif identified by ARTEM in all DNA-containing and (C) RNA-containing PDB entries. (D) A parallel all-adenine tetrad match identified by ARTEM at RMSD = 0.343 Å in a d[T_2_AG_3_] DNA repeat tetramer, PDB entry 2JWQ, chains A, B, C, D. (E) Top-view and (F) side-view of a false all-guanine match spanning residues from two adjacent non-parallel tetrads, identified by ARTEM at RMSD = 1.202 Å in an RNA iMango-III aptamer, PDB entry 6E8S, chain B.

**Supplementary Figure S5.**
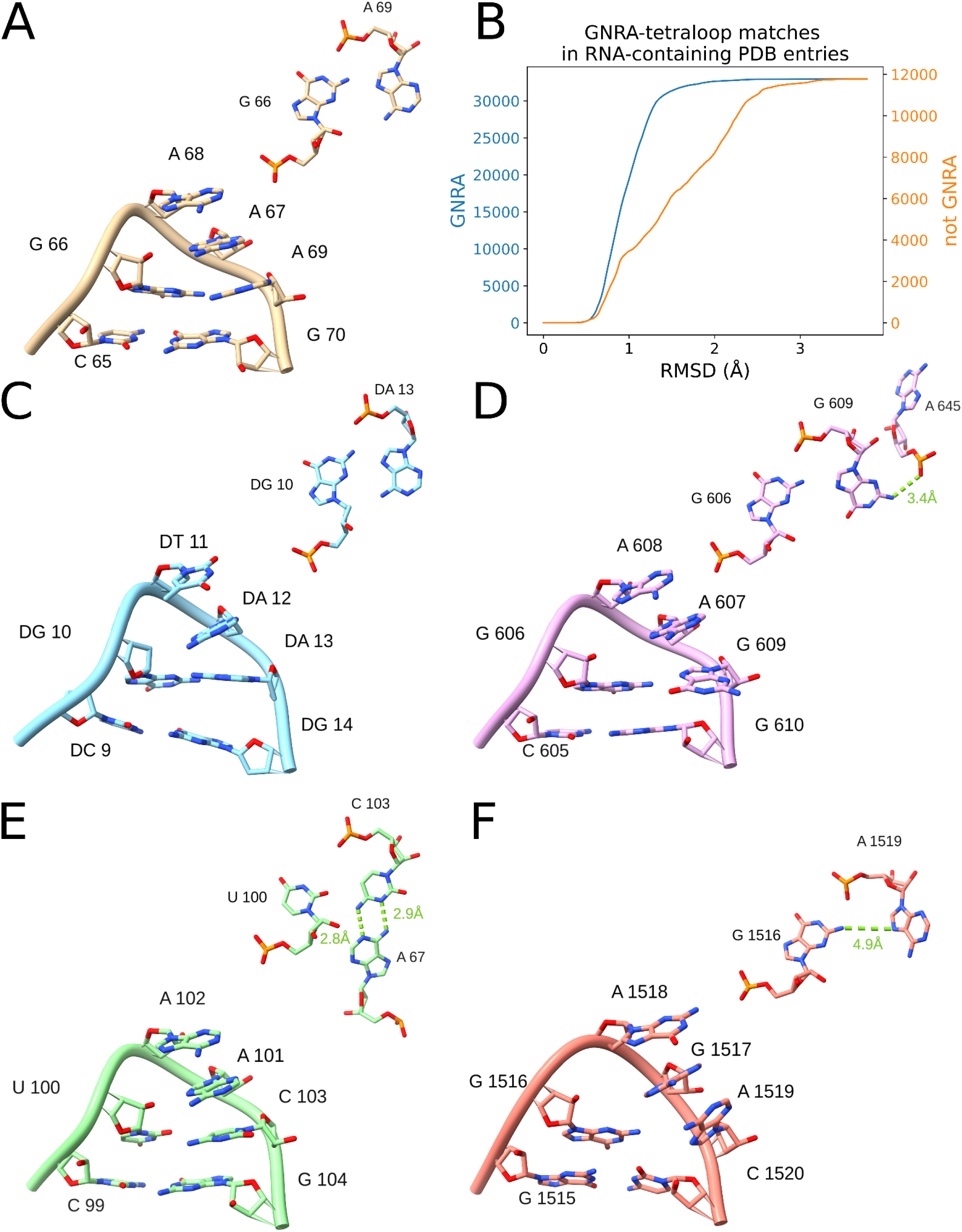
ARTEM search for the GNRA-tetraloop motif. (A) Reference GNRA-tetraloop instance from a THF riboswitch, PDB entry 3SUX, chain X. (B) GNRA (in blue) and non-GNRA (in orange) matches of the motif identified by ARTEM in all RNA-containing PDB entries. (C) GNRA match identified by ARTEM at RMSD = 0.986 Å in a dumbbell DNA, PDB entry 2N8A, chain B. (D) GAAG match identified by ARTEM at RMSD = 0.749 Å in 18S rRNA, PDB entry 7QEP, chain 3. (E) UAAC match identified by ARTEM at RMSD = 0.99 Å in a lariat capping ribozyme, PDB entry 4P8Z, chain A. (F) GNRA match with a distorted tSH G-A base pair identified by ARTEM at RMSD = 1.51 Å in 16S rRNA, PDB entry 6DNC, chain A.

**Supplementary Figure S6.**
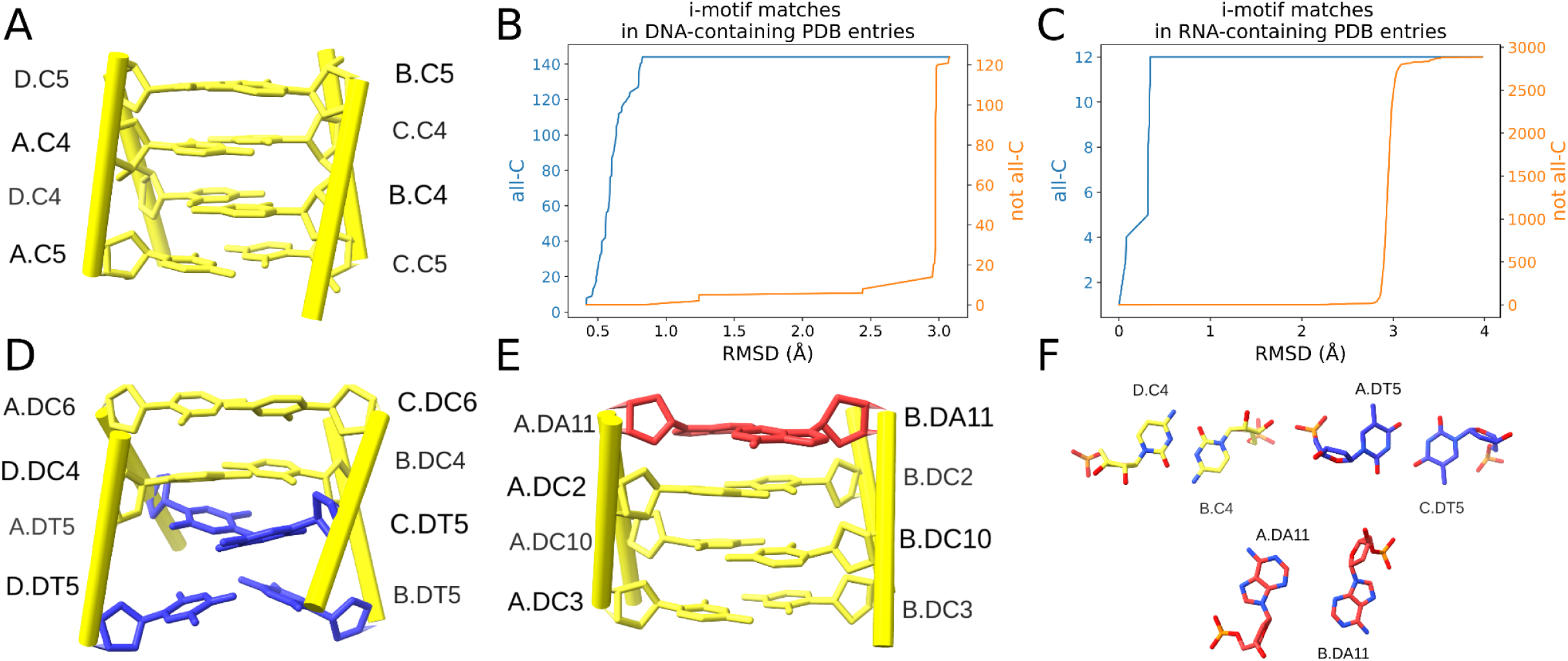
ARTEM search for the i-motif. (A) Reference RNA i-motif instance, PDB entry 1I9K. (B) all-cytidine (in blue) and not all-cytidine (in orange) matches of the motif identified by ARTEM in all DNA-containing and (C) RNA-containing PDB entries. (D) Thymine-containing match identified by ARTEM at RMSD = 1.004 Å in a DNA i-motif tetramer, PDB entry 2KKK. (E) Adenine-containing match identified by ARTEM at RMSD = 1.24 Å in a double hairpin DNA, PDB entry 1C11. (E) Top view of the tWW C-C, tWW T-T, and tSS A-A base pairs from the identified motifs.

**Supplementary Figure S7.**
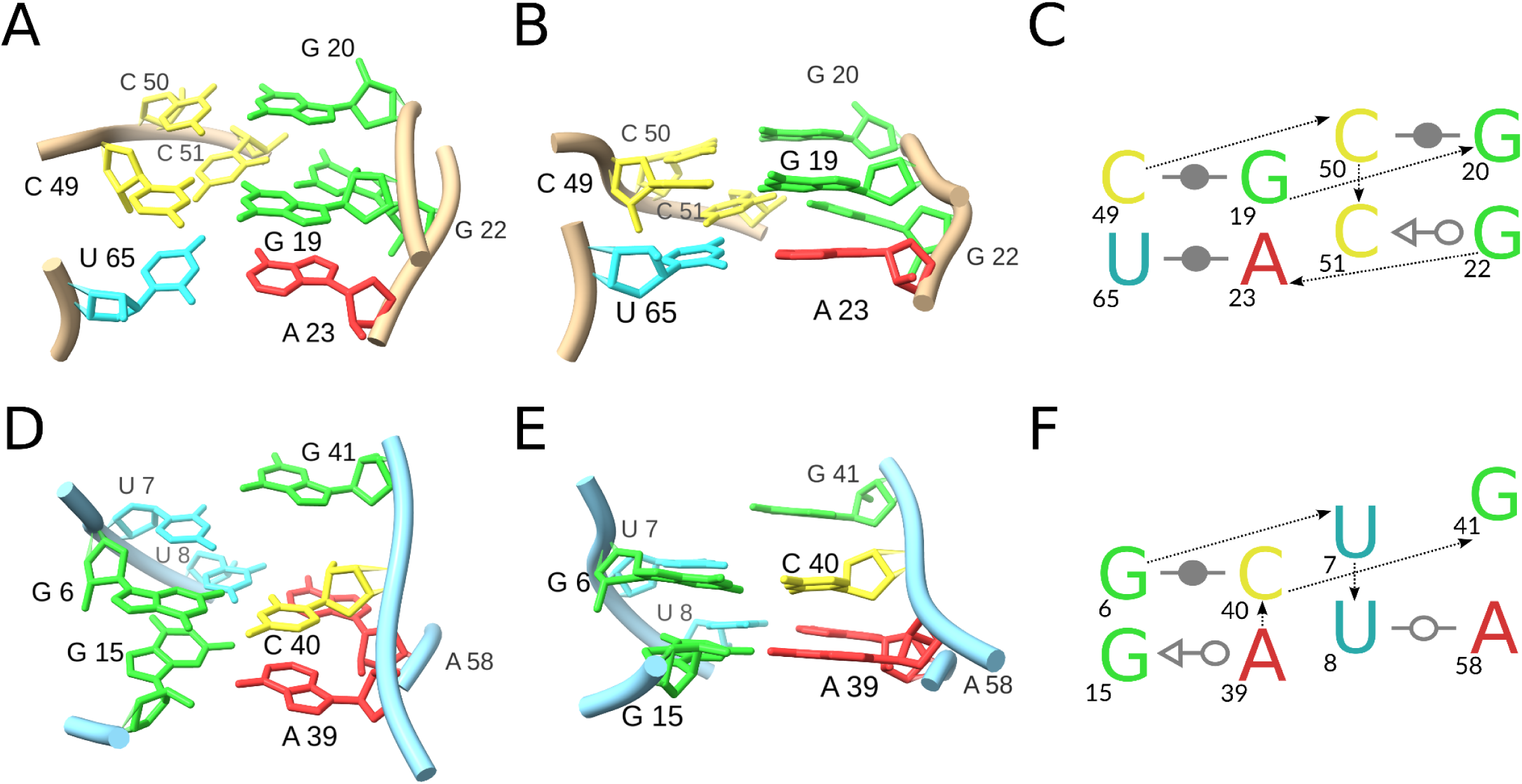
ARTEM search for the parallel-pairing motif. (A) top view, (B) side view, and (C) interaction scheme of the reference parallel-pairing motif from cap-independent translation enhancers from Pea enation mosaic virus RNA 2, PDB entry 8SH5, chain R. (D) top view, (E) side view, and (F) interaction scheme of the parallel-pairing motif match identified by ARTEM at RMSD = 1.679 Å in a NAD+ -II riboswitch, PDB entry 8GXC, chain A.

**Supplementary Table S1.** Kink-turn characteristics of representative group II intron structures.

| PDB entry chain | 3IGI chain A | 5G2X chain A | 6CHR chain A | 6ME0 chain A | 7UIN chain B | 8H2H chain A | 8T2S chain B |
| --- | --- | --- | --- | --- | --- | --- | --- |
| Intron | group IIC | group IIA | group IIB | group IIC | group IIC | group IIA | group IIC |
| Coordination loop | C148-C157<br>G220-G226 | G203-U210<br>A339-C348 | A186-C197<br>G293-U299 | A243-C253<br>G362-U368 | C151-C160<br>G228-G234 | G203-U210<br>A339-C348 | C151-C160<br>G228-G234 |
| L1-1n base pair | U150-A224(syn) |  | G188-A297 | G245-A366 | U153-A232 |  | U153-A232 |
| $\kappa$ extension bulge | A134-A144 | A185-A201 | A171-A182 | A228-A239 | A137-A147 | A185-A201 | A137-A147 |
| $\kappa$ extension - coordination loop interactions | tHW<br>A138-U150(L1)<br>stack<br>A137-A151(L2) | stack<br>U195-A344<br>stack<br>U193-G207 | ribose-base<br>A175-U189<br>(L1.1) | N4-O6 H-bond<br>C232-G245 | tHW<br>A141-U153(L1)<br>stack<br>G140-A154(L2) | stack<br>U195-A344<br>stack<br>U193-G207 | tHW<br>A141-U153(L1)<br>stack<br>G140-A154(L2) |
| σ-$\sigma'$ BWE | | | | A771<br>A248(L3)<br>A769 | | | |
| $\delta'$-δ base pair | G155-C180 | | G195-C255 | A251-U326 | G158-C183 | | G158-C183 |
| EBS3 | A223 |  | G296 | C365 | A231 |  | A231 |
| IBS3 | A7, chain B |  | C623 | DG15, chain B | DT36, chain A |  | G1 |
| Coordination loop residues exposed to reverse transcriptase |  |  |  | C247(L2)<br>G249(1b)<br>C365(EBS3) | U155(L3)<br>A231(EBS3) |  | U155(L3)<br>A231(EBS3) |
| DIII kink-turn |  |  |  |  | G343-U352<br>A366-C372 |  | G343-U352<br>A366-C372 |
| Coordination loop kink-turn was identified by | ARTEM $\geq 12$nt<br>ARTEM $\geq 11$nt | | ARTEM $\geq 11$nt | ARTEM $\geq 12$nt<br>ARTEM $\geq 11$nt<br>RNAMotifScanX | ARTEM $\geq 12$nt<br>ARTEM $\geq 11$nt | | ARTEM $\geq 12$nt<br>ARTEM $\geq 11$nt |
| DIII kink-turn was identified by | | | | | ARTEM $\geq 11$nt<br>DSSR<br>RNAMotifScanX | | DSSR<br>RNAMotifScanX |

